# Non-invasive *in vivo* lactate monitoring via NIR spectroscopy

**DOI:** 10.64898/2026.06.23.733906

**Authors:** Tibor Lehnert, Sebastian Seidel, Jonas Euchner, Adrian Thierbach, Felix Schmidt, C. Mohan Ögün, Wilfried Hermes

**Author notes:** **Corresponding Author**: Tibor Lehnert - trinamiX GmbH, Industriestraße 35, 67063 Ludwigshafen, Germany;, Celal Mohan Ögün - trinamiX GmbH, Industriestraße 35, 67063 Ludwigshafen, Germany;, General.

## Abstract

We present a non-invasive approach for continuous monitoring of lactate dynamics *in-vivo* using near-infrared (NIR) spectroscopy. Lactate-related spectral features were measured non-invasively within the overtone region (1600–1850 nm). Several anatomical measurement sites were evaluated, and the middle phalanx of the dorsal finger emerged as the most promising location due to its superior spectral quality and stable tissue perfusion, becoming the exclusive site for all further experiments. Across multiple exercise sessions, predictive models achieved high within-day accuracy (R^2^ *≥* 0.8), while cross-day performance was affected by spectral drift and physiological variability. A dynamic offset-correction procedure effectively mitigated these baseline shifts, enabling stable prediction accuracy across days, weeks, and subjects. These findings demonstrate the feasibility of NIR-based lactate estimation and highlight the importance of adaptive correction strategies for reliable long-term, non-invasive monitoring.

## Introduction

Lactate is a key metabolite of anaerobic glycolysis and is widely recognized as a biomarker for tissue hypoxia, hemodynamic instability, and metabolic dysregulation [1, 2]. In critical care, hyperlactatemia is characteristic of sepsis, septic shock, myocardial infarction, and respiratory failure. Numerous studies confirm its prognostic relevance: elevated lactate levels correlate with increased mortality, and lactate clearance within the first 24 hours serves as a dynamic parameter for guiding resuscitation [1, 3]. In sports medicine, lactate profiling is essential for determining the aerobic–anaerobic threshold and optimizing training strategies [2]. The current gold standard for lactate assessment is invasive blood sampling, either via laboratory analysis or point-of-care devices. While handheld analysers provide acceptable accuracy within clinically relevant ranges, they require repeated blood draws [2]. This approach imposes procedural burden, increases resource utilization, and creates informational gaps, as measurements are typically performed at multi-hour intervals. Continuous trend monitoring — critical for therapeutic decision-making — remains impractical with this methodology. These limitations underscore the need for alternative strategies enabling continuous, non-invasive, patient-centric monitoring [3].

Recent research focuses on three biofluids: sweat, saliva, and interstitial fluid (ISF). Sweat-based sensors allow epidermal, real-time monitoring but face challenges such as variable sweat rates and complex correlation with blood lactate [3–5]. Saliva offers convenient sampling yet suffers from matrix variability and flow inconsistencies [4]. ISF is considered physiologically closer to blood; approaches include reverse iontophoresis and minimally invasive microneedle systems [3, 5]. A recent review emphasizes that ISF-based sensors, particularly microneedle arrays, have demonstrated promising accuracy and short lag times compared to venous lactate, but require robust calibration and validation under dynamic conditions [3].

Optical spectroscopy — particularly near-infrared (NIR) and short-wave infrared (SWIR) methods (wavelengths range 1300 nm - 2500 nm) — has emerged as a promising approach for label-free lactate detection [6, 7]. Foundational work by Budidha et al. demonstrated the feasibility of lactate quantification using UV/Vis–NIR–MIR spectroscopy combined with multivariate calibration using partial least squares regression (PLSR), achieving high predictive accuracy in controlled solutions and identifying key absorption bands critical for future *in vivo* applications [8]. Complementary modelling studies emphasize the importance of optical path length, tissue scattering, and water absorption as major determinants of transcutaneous sensing performance [6]. Translating these principles into clinical practice, De Ridder et al. reported a first-in-human feasibility study of an implantable NIR spectroscopy sensor capable of multi-analyte monitoring—including lactate—over 28 days, achieving clinically relevant accuracy and demonstrating the potential for long-term, continuous metabolic surveillance [9].

Evidence from *in vitro* and *in vivo* studies further supports the viability of NIR spectroscopy for lactate measurement. Lafrance et al. showed *ex vivo* studies that NIR combined with chemometric modelling such as PLSR can quantify lactate in undiluted plasma across exercise-induced ranges with excellent agreement to enzymatic reference methods (R^2^*≈* 0.995; RMSCV *≈* 0.51 mmol/L) within the 2050–2400 nm spectral window [10]. Extending this concept to whole blood, Lafrance et al. demonstrated that near-infrared transmission spectroscopy can accurately measure lactate in human blood samples, confirming the robustness of NIR-based calibration models under physiologically relevant conditions [11].

At the design level, *in silico* investigations using Monte Carlo simulations have clarified constraints for SWIR-based transcutaneous sensing: photon penetration is shallow (*≈*1.3 mm), detected power is low (*≈* 0.3–2.5% of incident), and optimal configurations centre around 1684 nm with *≈* 1 mm source–detector spacing—necessitating highly sensitive detectors and robust calibration strategies [12]. Finally, keratinized sites such as the nail plate offer unique advantages for optical sensing due to their low water content compared to skin. Infrared studies of nails demonstrate diagnostically useful spectral windows and sampling geometries (attenuated total reflection, diffuse reflectance and photo acoustic spectroscopy) that primarily probe nail strata rather than the perfused nail bed, positioning nails as attractive portals for non-invasive spectroscopy [13].

The study by Sowa et al. provided the foundation for subsequent *in vivo* lactate measurements using NIR diffuse reflectance spectroscopy. It demonstrated that NIR can reliably detect lactate changes greater than 2 mmol/L (clinically relevant), indicating that non-invasive lactate monitoring is feasible—particularly when individual reference spectra are used [14].

The Surviving Sepsis Campaign Guidelines (2021) emphasize lactate as a marker of hypoxia and prognosis [1]. Lactate is integral to the Sepsis-3 definition of septic shock and is recommended for both screening and therapeutic targeting. Guidelines advocate serial or continuous lactate measurements to capture dynamic trends. In practice, however, monitoring is predominantly invasive and intermittent, resulting in sporadic data, patient discomfort, and high resource demand [1]. This creates a compelling clinical rationale for non-invasive, continuous lactate monitoring: such systems could facilitate guideline adherence, enhance patient safety, and improve therapeutic precision [3].

Non-invasive technologies promise high-frequency time-series data essential for sepsis management and athletic performance optimization. They minimize pain, infection risk, and iatrogenic anemia, reduce logistical complexity, and enable integration into telemedicine frameworks [4]. Despite these advantages, challenges persist, including reliable correlation with blood lactate, signal integrity in tissue, and long-term sensor stability. The development of robust, biocompatible materials and rigorous clinical validation will be pivotal for translating these innovations into routine practice [3, 5, 6, 9].

## Results and discussion

Across a series of *in vitro* and *in vivo* studies, we established and iteratively refined a NIR spectroscopy approach for the non-invasive estimation of lactate dynamics during and after physical exercise. The collective findings consistently demonstrate the feasibility of capturing physiologically relevant changes in lactate concentration *in vivo*, while also highlighting several methodological constraints and sources of variability that have been incorporated into the development of an optimized measurement and modeling strategy.

The use of *in vitro* measurements was necessary from an application-oriented perspective: *In vivo*, the permissible irradiance is strongly limited (irradiance limit: 3556 W/m^2^ for 10 s exposure time [15]), and selecting an appropriate optical window allows the measurement system to be designed such that the available photons effectively fill the detector’s dynamic range. This ensures that even under clinically constrained illumination conditions, the acquired signal remains within a usable and quantifiable range, thereby improving robustness and sensitivity of lactate detection in practice. Therefore, it is crucial to examine the spectral regions that offer the most favorable conditions for such measurements. Within the SWIR spectral range (1100–2500 nm), four distinct regions are of particular relevance due to the pronounced variability in the vibrational modes of organic molecules and the substantial penetration depth of light in biological tissues. These regions, designated as therapeutic windows and enumerated using Roman numerals (I–IV), are defined as follows: Window I spans 650–950 nm, Window II spans 1100–1350 nm, Window III spans 1600–1870 nm, and Window IV spans 2100–2300 nm [16].

Our infrared spectra were recorded in the range of window III (cf. fig. 1 (a)) instead of window I and II because of reduced photon scattering and minimal interference from dominant chromophores such as hemoglobin and melanin [17] resulting in enhanced detectability of lactate-specific absorption features, which were reported, according to *in-silico* simulations, to be at 1684 nm [12]. Window IV is also not considered due to the higher noise level and poor repeatability compared to spectral window III.

**Figure 1:**
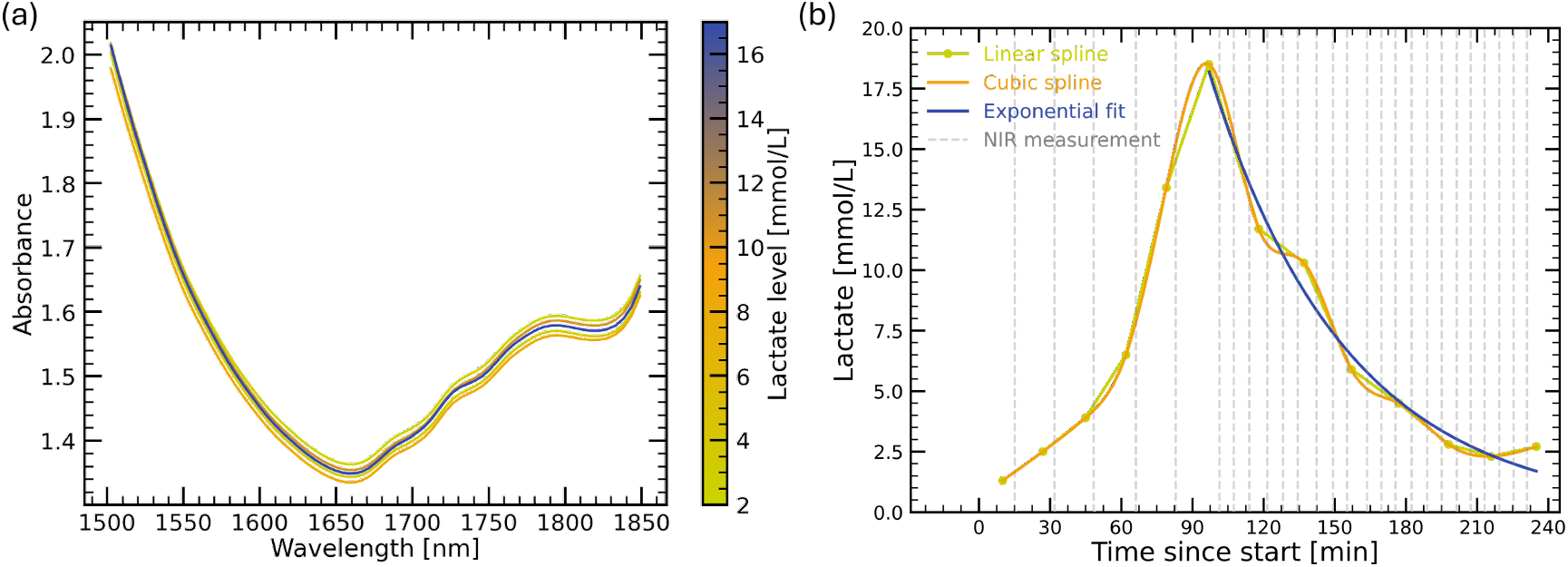
(a) Raw IR spectra acquired during the measurements in Window III. The color coding represents different lactate concentrations associated with each spectrum. No clear trend or systematic variation in the spectral features with respect to lactate levels can be observed without further data processing and analysis. (b) Interpolated reference lactate values are determined because NIR measurements and capillary blood sampling were not temporally aligned. A cubic spline was used to reconstruct the full lactate time course across both the build-up and decay phases, while an exponential fit was applied specifically to model the decay region. A linear spline served as an initial approximation. The gray dashed vertical lines indicate the times at which NIR measurements were taken. Interpolated reference values are obtained from the intersect of NIR measurements and the fit functions.

First *in vivo* measurements were conducted with two volunteers during high-intensity interval exercise, where blood lactate values reached up to approximately 20 mmol/L.

Figure 1 (a) presents the raw IR spectra obtained during the measurements. The different colors indicate varying lactate concentrations associated with each spectrum. At this stage, however, no clear trend or systematic relationship between spectral features and lactate levels is apparent without further analysis. This highlights the need for additional processing steps and careful alignment with reference measurements. However, a central challenge across all *in vivo* studies was the lack of exact temporal alignment between NIR scans and capillary blood sampling. To address this mismatch, reference lactate time courses were reconstructed using several interpolation strategies. A linear spline was initially employed as a first-pass approximation to estimate intermediate points. For the final reference reconstruction, a cubic spline was applied across the entire time range to model both the build-up and decay phases, while an exponential fit was used specifically to capture the kinetics of the elimination phase. Figure 1 (b) show these interpolations exemplary. The gray dashed vertical lines mark the time points at which the NIR measurements were recorded. The interpolated reference values are determined from the intersections between the NIR measurements and the fitted functions. Overall, the model predictions worked better based on the exponential decay interpolations.

Initial model evaluation was performed using a leave-one-out cross-validation (LOOCV) approach, in which prediction and reference values were compared separately for two subjects across three measurement days each. The resulting regression plots, which are shown in figure 2, demonstrated consistently robust performance, with linear fits (blue solid line) showing narrow confidence intervals, indicated by the black dashed lines. All models achieving coefficients of determination of R^2^ *≥* 0.8. This indicates that, under conditions where training and testing data originate from the same measurement day, the models are capable of capturing inter-individual lactate dynamics with high accuracy, given by the Root Mean Squared Error (RMSE), which is for all predictions below 1.85 mmol/L.

**Figure 2:**
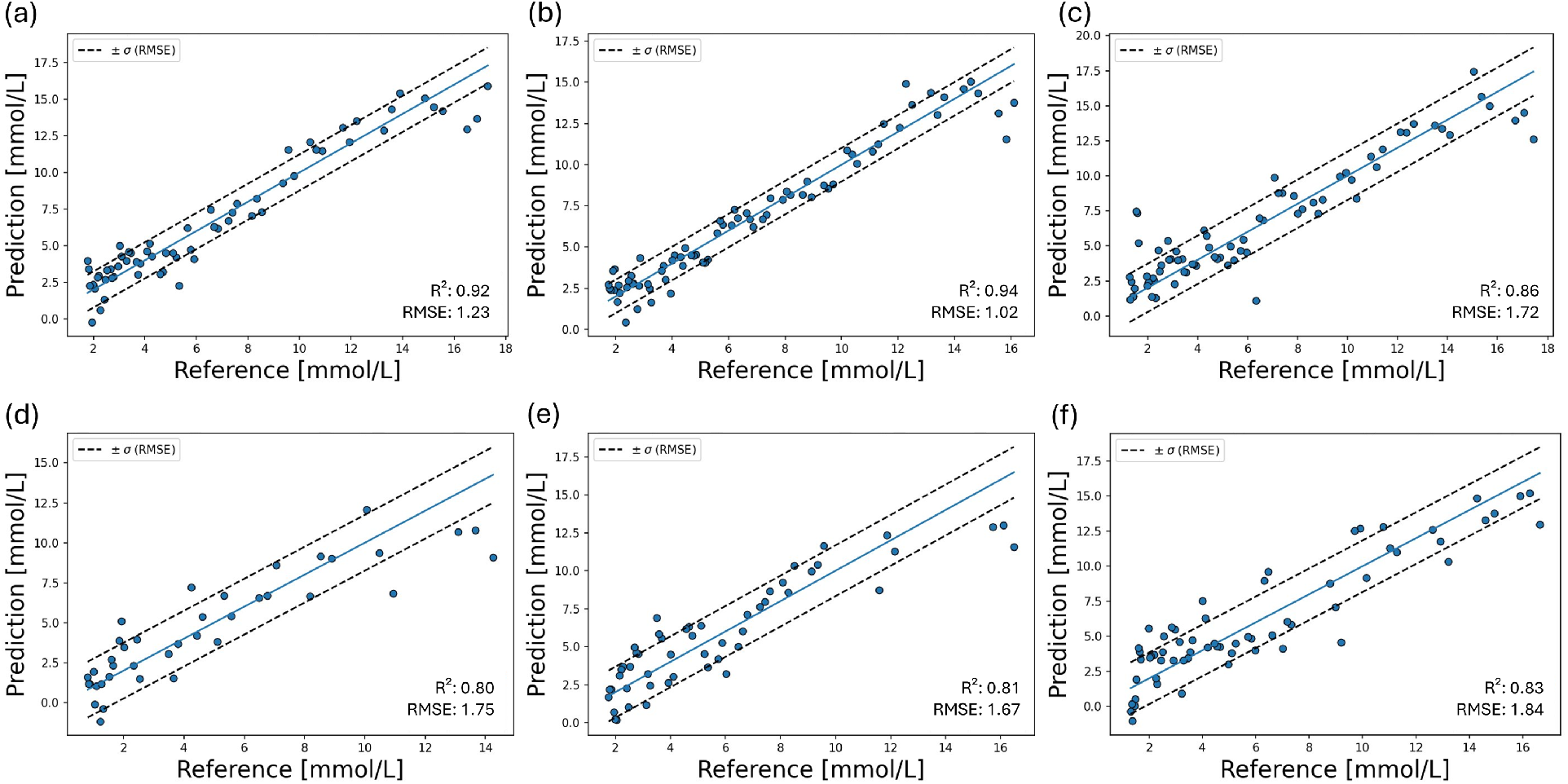
Lactate prediction based on LOOCV approach for the lactate clearance region, compared to reference measurements for two subjects across six measurement days: panels (a)–(c) show three different days for subject 1, and panels (d)–(f) three different days for subject 2. The blue line represents the linear fit, and the black dashed lines indicate the corresponding confidence interval. Reference values were obtained from capillary blood samples. For each plot, RMSE and R^2^ are reported; all regressions exhibit an R^2^ *≥* 0.8.

To evaluate the temporal robustness of the model and its ability to generalize beyond same-day training conditions, we examined cross-day model performance. In this analysis, the model was trained on data from one of the three measurement days and then used to predict lactate concentrations on all days. The results are plotted in figure 3 and show clear differences depending on which day served as the prediction target. Training on day three and predicting days one to three yielded the best generalization performance (R^2^ = 0.80), indicating stable behavior across days. Training on day one, and predicting all days, showed slightly reduced but still robust performance (R^2^ = 0.75). In contrast, training on day two, and predicting days one to three, resulted in the weakest performance (R^2^ = 0.58). This reduction was largely driven by three severe outliers originating from day three that produced negative lactate values in the training (cf. figure 3 (b)). Excluding these outliers, the remaining predictions followed a trajectory comparable to the other training conditions, suggesting that the lower overall R^2^ reflects isolated anomalies rather than a systematic failure to generalize. Although cross-day predictions show reduced accuracy compared to within-day LOOCV results, their RMSE remains in an acceptable range of approximately 2-3 mmol/L, indicating that predictive performance is still adequate for practical use despite increased temporal variability.

**Figure 3:**
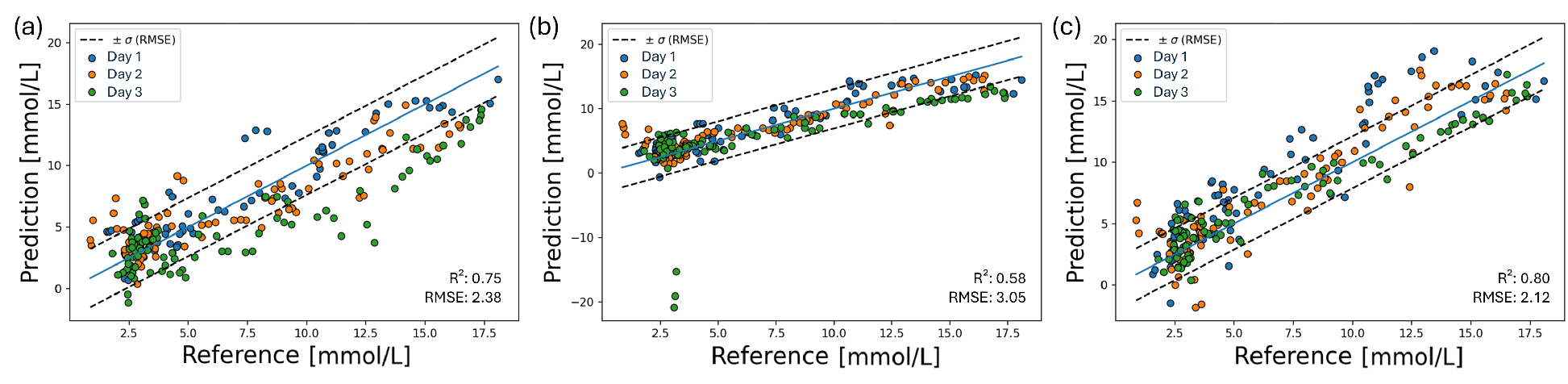
Cross-day model performance when training on a single day and predicting lactate concentrations across all days. Panel (a) shows results when training on day one and predicting days one to three. Panel (b) shows results when training on day two and predicting days one to three. Panel (c) shows results when training on day three and predicting days one to three. The best performance is achieved when training on day three (R^2^ = 0.80). Training on day one yields slightly lower but still robust performance (R^2^ = 0.75). The weakest performance occurs when training on day two (R^2^ = 0.58), primarily due to three strong outlier predictions with negative lactate values; excluding these outliers, the model’s behavior appears consistent and promising.

Taken together, the LOOCV and cross-day analyses highlight both the promise and the challenges of non-invasive lactate prediction. While within-day prediction performance is consistently high, cross-day generalization depends strongly on the characteristics of the training data. Importantly, the findings suggest that selecting training datasets with stable measurement conditions - such as those observed on day three - can substantially improve between-day prediction performance.

Combining multiple days for the training introduced distributional shifts that reduced prediction accuracy. This day-to-day variability was further observed as spectral drift, which created a systematic bias when applying models across days. Increasing the training set to seven separate measurement days reduced this bias, albeit at the cost of nominal predictive performance. Additional hardware modifications led to a substantial offset in recorded spectra; this was mitigated through a dynamic offset-correction procedure, which will be discussed in detail further below.

Building upon the cross-day evaluation, we next investigated whether model robustness could be further improved by substantially expanding the temporal diversity of the training dataset. For this purpose, seven measurement days from subject 1 were combined to form a consolidated training set for a predictive model, which was tested on another day. While this multi-day training strategy reduced day-specific spectral distortions, the predictions for subject 1 still exhibited a clear systematic offset relative to the reference values as shown in figure 4 (a). The large offset results in an negative R^2^ for the prediction (Fig. 4 (b)), indicating that the model is not suitable. The unpredictability can be observed via the large mismatch of the line of identity and the line of best fit shown in figure 4 (b). To address the offset-induced unpredictability, we implemented a dynamic offset-correction procedure, in which the average of all NIR measurements collected during the resting phase are shifted such that their predicted values align with the average of the corresponding capillary blood reference values, leading to a better overlap of all blood reference values and the NIR measurements in figure 4 (c). This alignment compensates for accumulated spectral drift and baseline shifts, significantly improving predictive accuracy (corrected values: 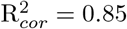 RMSE_*cor*_ = 1.61 mmol/L), so that a good overlap of the line of identity and the line of best fit appears (cf. Fig. 4 (d)). The predictive model developed, which also includes dynamic offset correction, shows good stability even after a longer period of time. Test subject 1 was measured again 1 month and 2 months later, and the prediction shows good values with R^2^ = 0.86 after one month (Fig. 4 (e)) and R^2^ = 0.76 after two month (Fig. 4 (f)).

**Figure 4:**
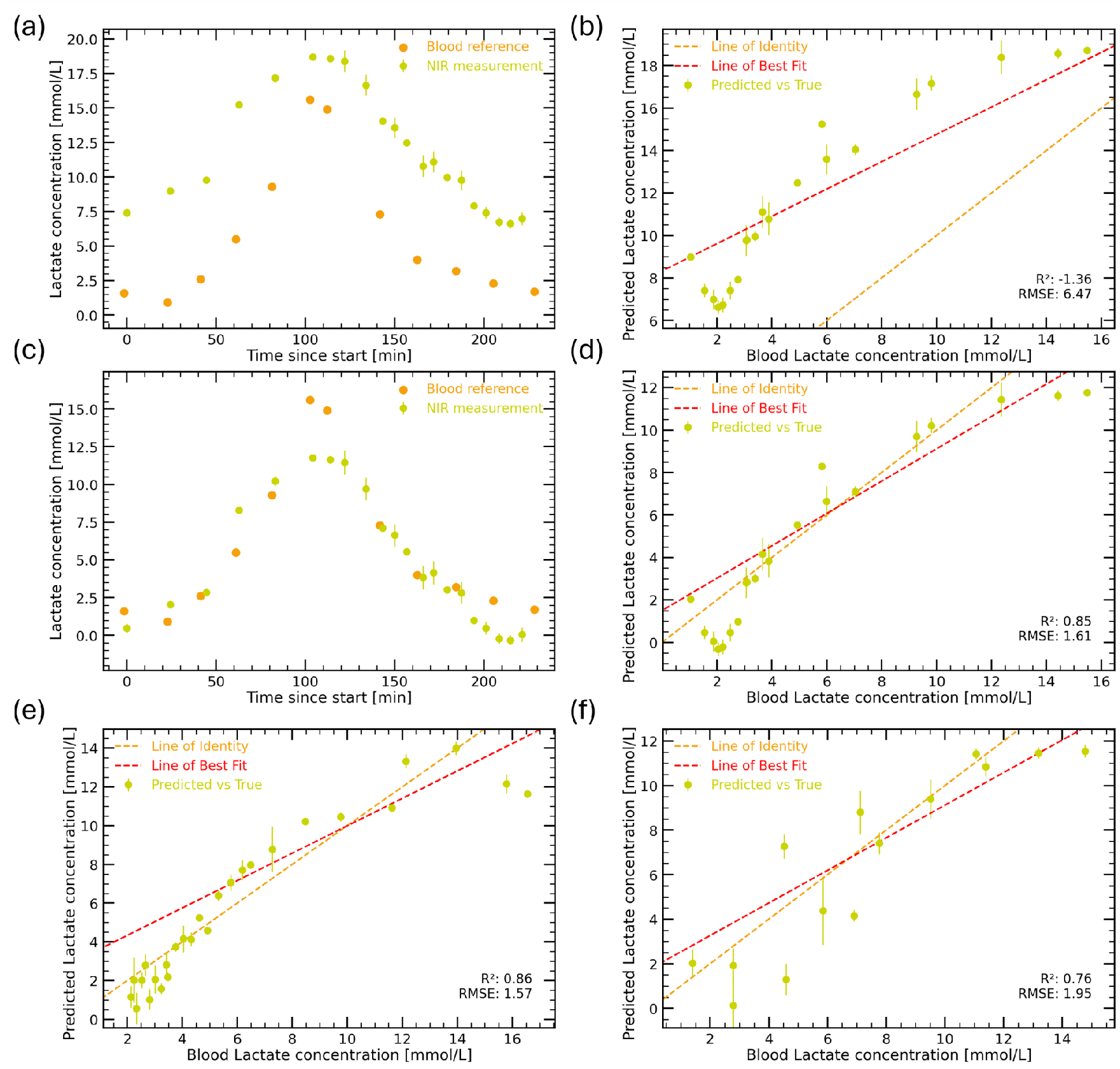
Prediction performance of the developed lactate-prediction model for different exercise days of subject 1. Panel (a) shows NIR data recorded two weeks after model training, exhibiting a clear offset between blood reference and NIR measurements. Consequently, the corresponding prediction in panel (b) shows poor performance with a negative R^2^ value (R2 = −1.36, RMSE = 6.47 mmol/L), which is reflected by the lack of overlap between the line of identity and the line of best fit. Panel (c) illustrates the effect of the implemented dynamic offset-correction procedure, leading to improved alignment between NIR and blood reference measurements. This correction results in a markedly improved prediction, shown in panel (d), with R^2^ = 0.85 and RMSE = 1.61 mmol/L, where the line of identity and line of best fit lie close together and exhibit nearly parallel slopes. Panels (e) and (f) display model performance with offset correction for data recorded one month and two months after the training period, respectively (R^2^ = 0.86, RMSE = 1.57 mmol/L; R^2^ = 0.76, RMSE = 1.95 mmol/L).

The slightly weaker performance observed two months after the training period (R^2^=0.76 instead of R^2^ *≈* 0.85, cf. figure 4) can be attributed to accumulated day-to-day variability in measurement conditions, such as changes in skin hydration or perfusion, which alter the spectral response. Occasional measurement artifacts or outliers further influence the regression metrics, especially when sample sizes per session are limited. Additionally, the inherent analytical uncertainty of capillary blood lactate measurements contributes to increased variability, particularly at low concentrations, where handheld analyzers show broader prediction intervals and systematic biases compared to reference laboratory methods [23, 24].

To assess inter-subject generalizability, the lactate-prediction model, which was trained on subject 1, was applied to subject 2. All available data from Subject 2 were compiled for this purpose. Without the offset correction, a negative R^2^ = *−* 5.87 was determined, indicating that the model is not valid, shown in figure 5 (a). However, the predictions exhibit a noticeable offset and the slope of the linear fit closely matches the expected physiological trend, demonstrating that the underlying relationship between spectra and lactate concentration is preserved. The dynamic offset-correction substantially improves the accuracy and reduced residual prediction errors resulting in a positive 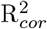 (corrected values: 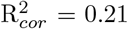, RMSE_*cor*_ = 3.93 mmol/L).

**Figure 5:**
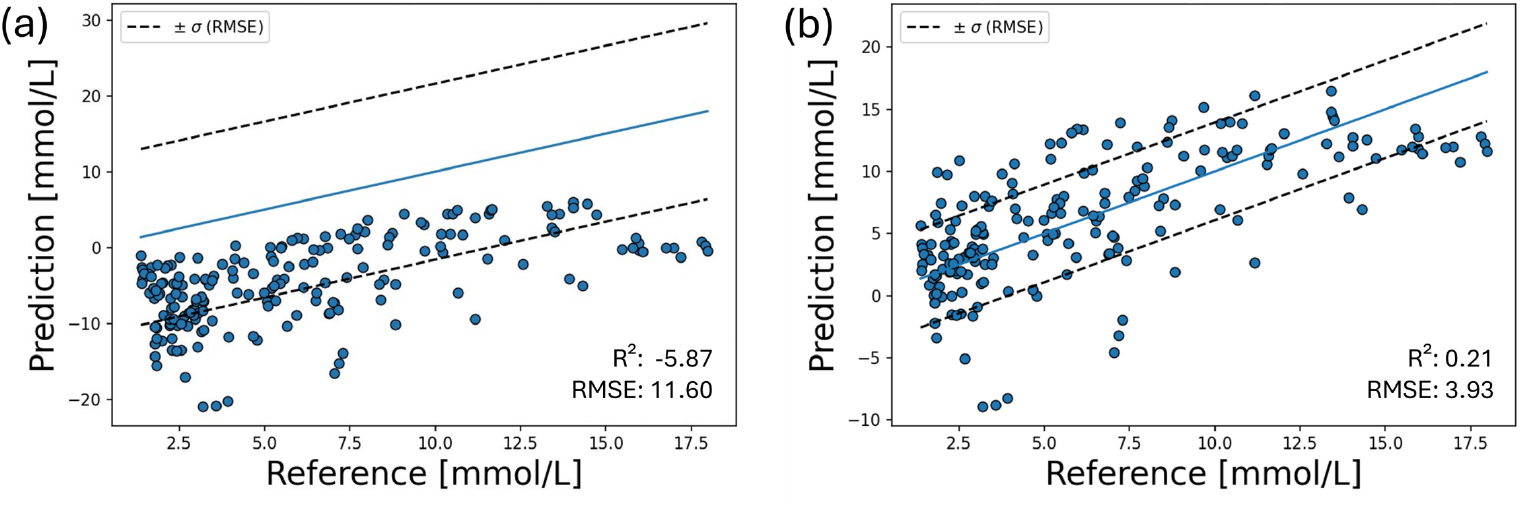
Prediction performance for subject 2 on all available measurement days, using the model trained on subject 1. Panel (a) reveals an offset between predicted and reference lactate values, similar to the inter-day offsets previously observed in subject 1. Despite this bias, the data points follow the same underlying trend as the model predictions. Applying the dynamic offset-correction procedure (b) substantially reduces both prediction error and systematic bias, resulting in improved accuracy with RMSE_*cor*_ = 3.93 mmol/L and coefficient of determination 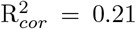.

Model generalization across participants is currently limited. Although applying the offset correction yielded a significant improvement, predictions for unseen subjects remain suboptimal. Collecting more data from a broader, more diverse participant pool is expected to strengthen the model and enhance cross-subject robustness. Encouragingly, this initial iteration indicates that both the model and its trajectory predictions are trending in the right direction, providing a solid basis for continued refinement.

Overall, this work demonstrates the feasibility of non-invasive lactate estimation using near-infrared (NIR) spectroscopy and highlights both the potential and the limitations of the approach. *In-vitro* experiments enabled the identification of robust lactate-sensitive regions, particularly around the overtone, which exhibited the highest repeatability and signal stability. These findings formed the foundation for all subsequent in-vivo measurements, which were conducted exclusively at the dorsal middle phalanx of the finger. The finger proved superior to the volar forearm for ISF-based NIR lactate monitoring, consistently delivering higher spectral quality and more accurate models—consistent with its dense peri-capillary network, and pronounced sudomotor/evaporative dynamics and faster ISF turnover at the dorsal finger.

Within single measurement days, prediction models achieved high accuracy (R^2^ *≥* 0.8; RMSE < 1.85 mmol/L), demonstrating that NIR spectra reliably capture lactate-related changes when measurement conditions remain stable. However, performance variations were observed due to spectral drift, physiological variability (e.g., changes in skin hydration, perfusion, or tissue scattering), and occasional measurement artifacts. These factors produced systematic offsets that limited the direct transferability of models over time. Combining multiple days into the training set reduced day-specific variability but introduced new distributional shifts, underscoring the sensitivity of linear regression models to baseline changes.

To address these temporal inconsistencies, a dynamic offset-correction procedure was developed. This method aligned the average predicted NIR values at rest with the corresponding capillary blood measurements, effectively compensating for day-to-day baseline shifts. Offset correction substantially improved prediction accuracy across repeated sessions, restoring high performance even one and two months after the training period. The improvement suggests that the primary limitation for cross-day generalization is a correctable offset rather than a change in the underlying spectral–physiological relationship.

Intersubject evaluation showed similar behavior: although absolute prediction offsets differed between individuals, the slope of predicted lactate dynamics was preserved. After offset correction, prediction errors decreased markedly, indicating that subject-specific differences can be largely compensated through minimal calibration rather than entirely new training datasets. This finding suggests that NIR-based lactate monitoring is transferable across users when supported by lightweight adaptive correction strategies.

Overall, the results confirm that NIR spectroscopy can non-invasively track lactate dynamics with high fidelity under controlled conditions and that the fundamental spectral–lactate relationship is stable across time and individuals. At the same time, the work highlights the need for robust drift-handling mechanisms to ensure long-term applicability. The dynamic offset-correction approach presented here represents a practical step toward more reliable daily use, enabling stable cross-day and cross-subject predictions without extensive retraining.

In conclusion, ISF-based NIR lactate monitoring is technically feasible and offers promising potential for continuous, non-invasive metabolic assessment. Future work should aim to minimize spectral drift through improved sensor hardware, explore adaptive or self-calibrating machine-learning methods, and validate the approach in larger and more diverse populations within clinical studies with, e.g, gas chromatography as reference analytics. These developments will be essential to transition NIR-based lactate estimation from proof-of-concept toward robust real-world application.

## Experimental

### NIR Setup

A custom-built Fourier-transform infrared (FTIR) spectrometer was utilized for near-infrared (NIR) measurements, specifically designed for biomarker detection in biological samples. The system employed a broadband light source covering a relevant range from 1500 nm to 2200 nm, coupled with a high-resolution interferometer and an InGaAs detector optimized for low-noise performance. The spectrometer achieved a signal-to-noise ratio (sSNR) of approximately 330,000 in the overtone region (1550–1800 nm) and at least 180,000 in the combination band region (up to 2200 nm). The sSNR was determined using a 99% reflectance standard from Labsphere for measuring the whole spectrum with an exposure time of 2 sec (corresponding to 4 scans). Spectral acquisition was performed with a resolution of 32 cm^*−*1^, and each measurement consisted of 120 scans to enhance signal quality. The optical setup was configured for spatial resolved reflectance measurements through the tissue at the dorsal side of the middle phalanx of the finger, using a custom-designed sample interface (cf. Fig. 6 (a)) to ensure reproducible positioning and minimize motion artifacts.

**Figure 6:**
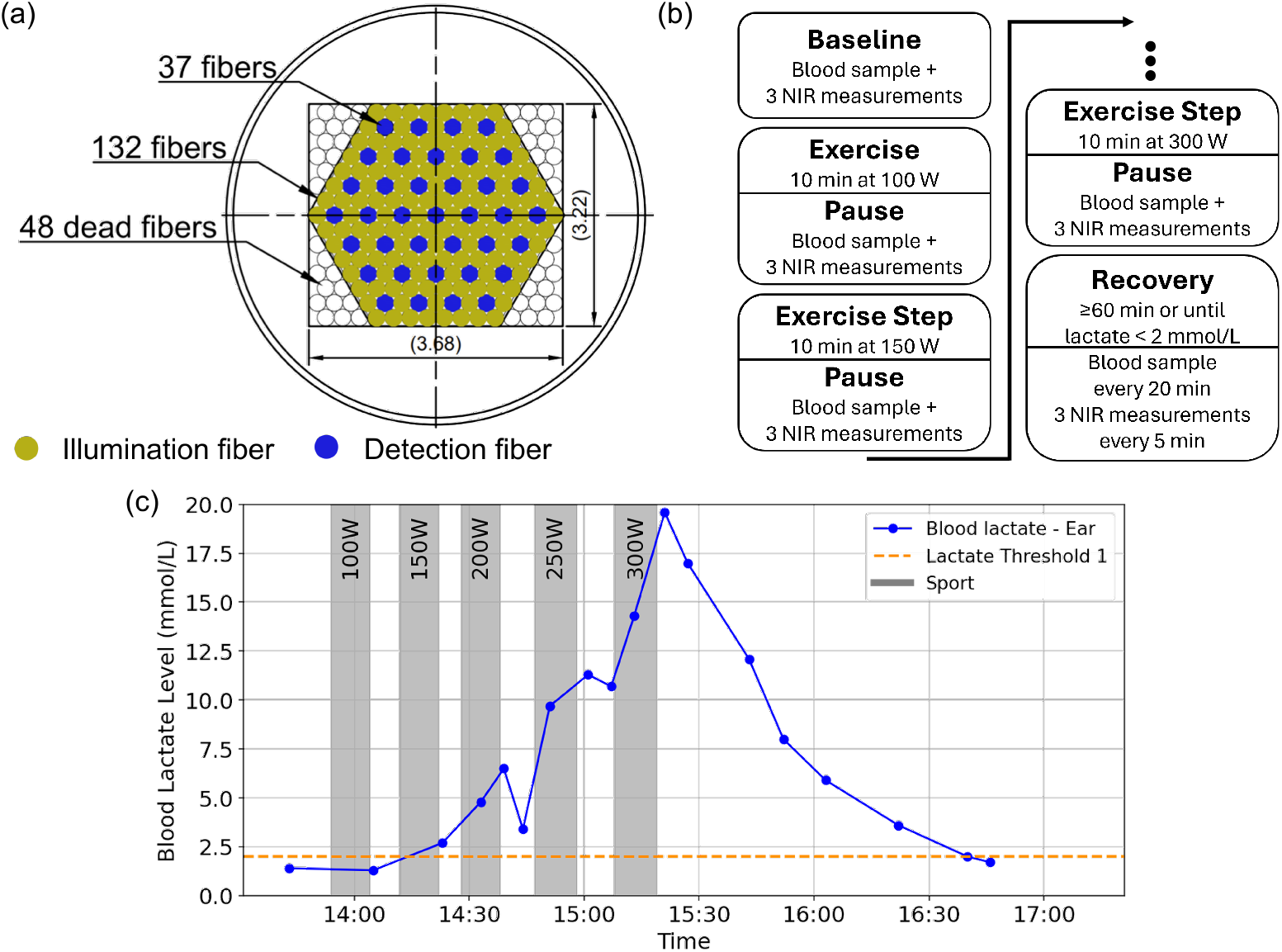
Cross-section of the specially designed fiber (a), featuring 37 fibers for detection (blue) and 132 fibers (green) for illumination with an overall size of 3.7 mm *×* 3.22 mm. 48 fibers were included to facilitate processing, but they serve no purpose later on (dead fibers). (b) shows the measurement plan which began with a baseline assessment comprising three NIR scans and one capillary blood sample for reference analysis. The exercise protocol started with 10 minutes of cycling at 100 W on an ergometer, followed by a 5-minute rest period during which three NIR scans and another capillary blood sample were collected. The workload was then increased in 50 W increments for subsequent 10-minute intervals, with the measurement sequence repeated after each stage until a maximum workload of 300 W was reached. Finally, recovery was monitored by acquiring three NIR scans every 5 minutes and a capillary blood sample every 20 minutes. (c) illustrates a real measurement procedure of a subject, where gray shaded regions denote the subject’s exercise periods and the blue curve represents the increase in blood lactate concentration measured via a capillary blood sample. The dashed line indicates the threshold below which lactate levels are considered within the normal range in the absence of physical activity.

The purpose of the custom-designed sample interface is to enable spatially separated illumination and readout from the target for light collection from a volume within the sample rather than only from its surface. The current design features a fiber bundle with a 200 *µ*m core diameter per fiber and a numerical aperture of 0.37. Germanium-doped glass is used to ensure high transmission in the near-infrared range. The bundle consists of 132 illumination fibers and 48 inactive fibers, with an overall size of 3.7 mm x 3.22 mm as shown in figure 6 (a).

### HIIT Session

Two subjects participated in high-intensity interval training (HIIT) sessions. Prior to exercise, baseline measurements were recorded, consisting of one blood sample and three NIR measurements to establish the initial state (cf. flowchart in Fig. 6 (b)). The exercise protocol began with 10 minutes of moderate cycling at 100 W on an ergometer, followed by a 5-minute pause during which three NIR measurements and one blood sample were collected. The measurement site was wiped with a dry cloth before NIR monitoring was started. The workload was then increased by 50 W for another 10-minute cycling interval, after which the same measurement routine was repeated. This stepwise increase continued until the subject reached 300 W. Following the workout, recovery was monitored for at least 60 minutes or until lactate levels returned to *<*2 mmol/L. During recovery, three NIR measurements were taken every 5 minutes and blood samples every 20 minutes. Each session lasted approximately 2 hours and 20 minutes. The procedure with lactate measurement data from capillary blood is illustrated in Figure 6 (c). The gray areas represent the subject’s exercise periods, while the blue curve indicates the increase in blood lactate concentration. During the exercise intervals, a clear increase of the lactate level is observed and a fast decrease during the recovery phase. For the presented results, two subjects were tested: subject 1 completed eleven sessions on different days, and subject 2 completed four sessions. Chemometric models were trained using data from subject 1 and validated on separate sessions of subject 1 as well as on data from subject 2.

### Lactate Measurements

The middle phalanx of the dorsal finger was used as the for measurements, since the literature identifies this position as high-dynamics site, characterized by pronounced sudomotor and evaporative activity. Key features include also high local sweat flux, elevated transepidermal water loss, and dense peri-glandular capillarization, all of which support faster local solute exchange and a high ISF-specific optical path [19, 20]. Lactate reference values were obtained using the Lactate Scout Sport device via capillary blood sampling. According to the manufacturer, the Lactate Scout Sport provides an accuracy of up to *±* 0.2 mmol/L for lactate concentrations between 0.5 to 6.7 mmol/L and within *±* 3% for values from 6.8 up to 25 mmol/L. It also offers a precision of 3-8 % depending on concentration, based on published technical specifications [25].

### Chemometry

Reference lactate concentrations obtained by capillary blood sampling were time-aligned to the NIR measurements by interpolation. Depending on the exercise phase, either cubic spline interpolation (covering build-up and decay phases) or exponential decay fitting (decay phase only) was applied. For each interpolation approach, partial least squares regression (PLSR) models were generated.

All spectra acquired during a measurement were cropped to the wavelength range of 1500–1850 nm. Subsequently, a grid search over commonly used spectral preprocessing steps was performed. The evaluated preprocessing combinations included spectral averaging, baseline correction, smoothing, derivative calculation, unit variance scaling, and scatter correction.

PLSR models were trained using either split cross-validation (CV) or leave-one-out cross-validation (LOOCV). The final model was trained using data from the first two measurement days of subject 1, while model selection was based on performance on the third measurement day of subject 1. The selected model was then further validated on an independent dataset obtained from subject 1 and subject 2.

## Author Information

### Author Contribution

‡T.L. and S.S. contributed equally to this work. The manuscript was written by T.L. and edited through contributions from all authors. S.S., J.E. carried out experiments. Data were analyzed by T.L., S.S., J.E., and A.M.T., reviewed and discussed by all authors. T.L., J.E., F.S., C.M.Ö., and W.H. planned and supervised the project. All authors have given approval to the final version of the manuscript.

## Notes

The authors declare the following competing financial interest(s): T.L., S.S., J.E., A.M.T., F.S., C.M.Ö., and W.H. are employed at trinamiX GmbH which commercializes spectroscopic sensor technologies. trinamiX GmbH and BASF have filed patent application(s) related to this work.

## Acknowledgements

The authors thank BASF SE for the financial support.

